# Cortico-subcortical functional connectivity profiles of resting-state networks in marmosets and humans

**DOI:** 10.1101/2020.07.14.202382

**Authors:** Yuki Hori, David J. Schaeffer, Atsushi Yoshida, Justine C. Cléry, Lauren K. Hayrynen, Joseph S. Gati, Ravi S. Menon, Stefan Everling

## Abstract

Understanding the similarity of cortico-subcortical networks topologies between humans and nonhuman primate species is critical to study the origin of network alternations underlying human neurological and neuropsychiatric diseases. The New World common marmoset (*Callithrix jacchus*) has become popular as a non-human primate model for human brain function. Most marmoset connectomic research, however, has exclusively focused on cortical areas, with connectivity to subcortical networks less extensively explored. In this study, we aimed to first isolate patterns of subcortical connectivity with cortical resting-state networks (RSNs) in awake marmosets using resting-state functional magnetic resonance imaging (RS-fMRI), then to compare these networks to those in humans using connectivity fingerprinting. While we could match several marmoset and human RSNs based on their functional fingerprints, we also found a few striking differences, for example strong functional connectivity of the default mode network with the superior colliculus in marmosets that was much weaker in humans. Together, these findings demonstrate that many of the core cortico-subcortical networks in humans are also present in marmosets, but that small, potentially functionally relevant differences exist.

## Introduction

The New World common marmoset (*Callithrix jacchus*) has become popular as a model for human brain function (Okano et al., 2016). Owing to a developed frontal cortex (Okano and Mitra, 2015) and the feasibility of creating transgenic marmosets (Park et al., 2016; Sasaki et al., 2009; Tomioka et al., 2017a; Tomioka et al., 2017b), the marmoset has become a promising candidate for assessing neuropsychiatric disorders, especially those involving frontal impairments that are more difficult to study in rodent models (Okano and Mitra, 2015). In the past few years, marmoset brain connectomics, including corticocortical anatomical connections (Majka et al., 2020; Majka et al., 2016), functional networks/connections (Hori et al., 2020a; Hori et al., 2020b), and white matter pathways (Liu et al., 2020; Schaeffer et al., 2017), are becoming increasingly well-studied. In addition, similarities of these connections have been found between marmosets and humans (Liu et al., 2020; Schaeffer et al., 2019a; Schaeffer et al., 2019b; Solomon and Rosa, 2014).

Demonstrating homologies across species is a challenging endeavor due to both limitations in measuring networks using the same method across species, and in identifying analogous brain areas to compare across vastly different brain morphologies. Resting-state functional magnetic resonance imaging (RS-fMRI) allows for circumvention of some of these challenges by allowing for non-invasive identification of robust and reproducible resting-state networks across different species (Damoiseaux et al., 2006; Fox and Raichle, 2007; Smith et al., 2013; Sporns, 2013). With recent advances in MRI hardware, we are now able to measure the functional networks/connectivities in awake marmosets (Belcher et al., 2013; Cléry et al., 2020; Hori et al., 2020b; Schaeffer et al., 2019c). Particularly, the marmoset’s small size is ideal for ultra-high field small-bore fMRI, affording high spatial resolution and signal-to-noise ratio (SNR) even in subcortical areas. Despite the ability to acquire MRI-based connectivity data in both marmosets and humans, the problem still stands of how to compare topologies amid major morphological differences. Connectivity fingerprinting have been offered as a method to circumvent this problem; this approach was originally proposed by Passingham and colleagues as a way to quantitatively evaluate the connections of a single cortical area with a selected set of other areas (Passingham et al., 2002). More recently, Mars and colleagues have suggested the feasibility of this approach as a tool for comparing various aspects of brain organization across and within species (Balsters et al., 2020; Mars et al., 2018; Mars et al., 2016; Schaeffer et al., 2020). Here, we employed this technique to compare cortico-subcortical fingerprints of resting-state networks (RSNs) in marmosets and humans, allowing for identification of inter-species similarities of cortico-subcortical connectivities.

We applied recent advances in hardware development for awake marmoset imaging, including a custom-made multi-array coil, a gradient coil (Handler et al., 2020), and an integrated head-fixation system (Schaeffer et al., 2019c) designed for small-bore ultra-high field MRI (9.4 T). This system allows for nearly motion-less, high spatial resolution, and signal-to-noise ratio (SNR) images. For human analyses, we used openly available datasets from the Human Connectome Project (HCP) (Van Essen et al., 2013). We used a data-driven approach via independent component analysis to identify RSNs in both marmosets and humans, then specified the subcortical connections with each cortical RSN. The cortico-subcortical functional fingerprints were created based on subcortical volumes of interests (VOIs), and were used to identify putative homologous RSNs between marmosets and humans.

## Results

### Cortico-subcortical RSNs in marmosets

The overarching objective of this study was to identify the subcortical areas related to each cortical RSN in marmosets, then to compare these cortico-subcortical connections between marmosets and humans. To do so, we first identified cortical functional networks in the marmosets (see Supplementary Fig. 1). After implementation of group ICA (Beckmann and Smith, 2004) using awake RS-fMRI data in only cortical regions, 6 components were identified as unstructured and/or physiological noise. The remaining 14 components demonstrated meaningful RSNs (Fig. 1). These RSNs were thresholded at z = 2.3 for visual purposes. Obtained RSNs were consistent with previously observed networks in marmosets (Belcher et al., 2013; Ghahremani et al., 2016; Hori et al., 2020b) such as default mode network (DMN) (Fig. 1A), attention network (ATN) (Fig. 1B), salience network (SAN) (Fig. 1C), left and right primary visual networks (pVIS-Lt/Rt) (Figs. 1D, 1E), orbitofrontal network (ORN) (Fig. 1F), high-order VISs (hVIS1-4: Fig. 1G-J), somatomotor networks ventral part (SMNv: Fig. 1K), dorsal part (SMNd: Fig. 1L) and medial part (SMNm: Fig. 1M), and premotor network (PMN: Fig. 1N).

**Figure 1.**
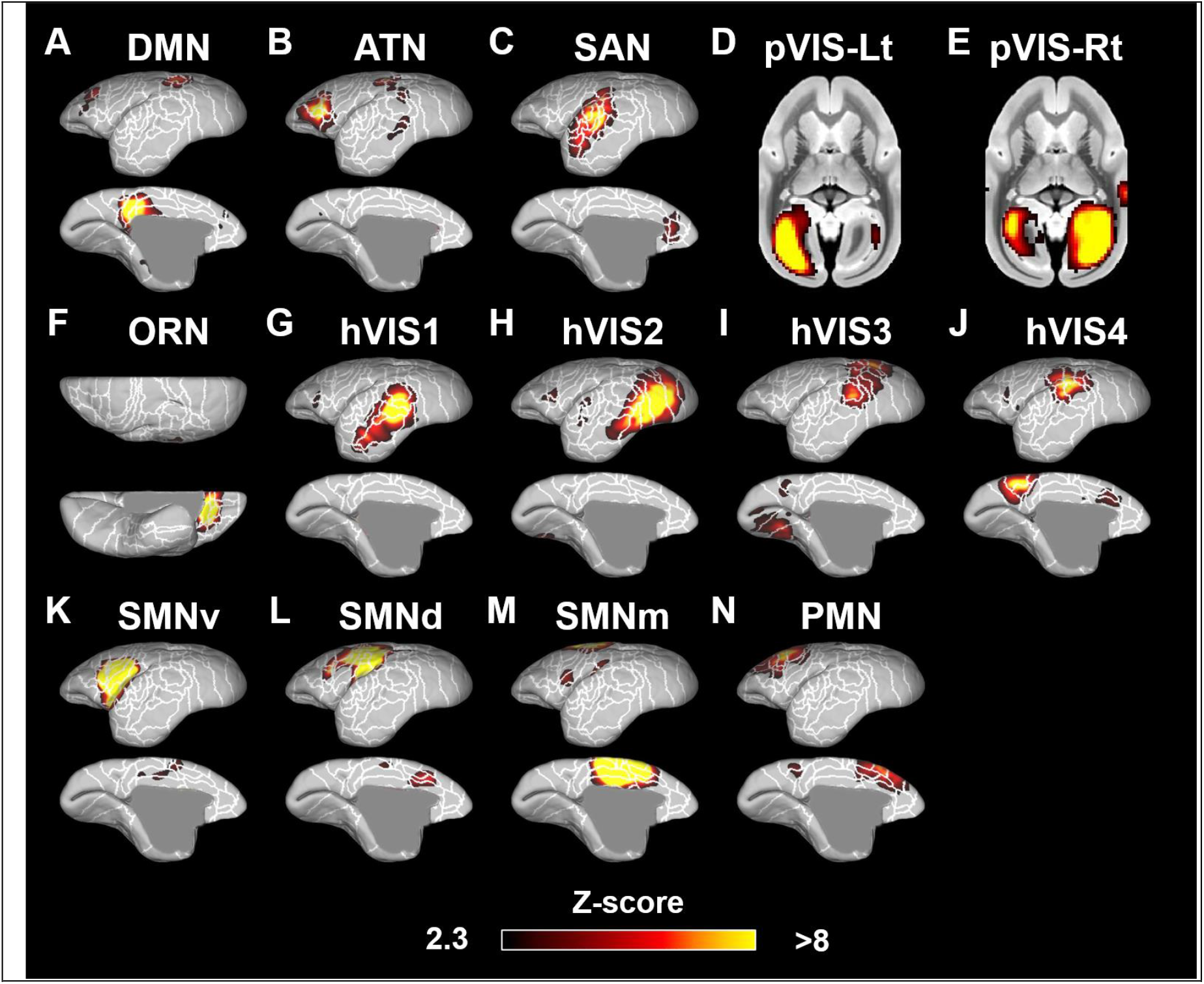
Fourteen components identified as resting-state networks in the marmosets. These networks were labeled based on previous studies (Belcher et al., 2013; Hori et al., 2020b) as follows: (A) default mode network (DMN); (B) attention network (ATN); (C) salience network (SAN); (D) left primary visual network (pVIS-Lt); (E) right primary VIS (pVIS-Rt); (F) orbitofrontal network (ORN); (G)-(J) high-order VIS (hVIS1-4); (K)-(M) somatomotor networks ventral (SMNv), dorsal (SMNd), and medial (SMNm); (N) premotor network (PMN). Color bar represents the z-score of these correlation patterns thresholding at 2.3. White lines show the cytoarchitectonic borders for reference (Liu et. al., 2018).

We calculated correlation coefficients between the time courses in each cortical RSN and the time courses in each subcortical voxel. The functional connectivity maps (z-score maps) in the subcortical areas were then averaged across scans. Averaged z-score values in each subcortical area corresponding to each RSN are shown in Supplementary Fig 2, and representative activation maps (z-score maps) are presented in Fig. 2. These z-score maps were normalized to be maximum z-value equal to 1 and were thresholded at 0.2 for visual purposes. The main subcortical area in the DMN (corresponding to Fig. 1A) was the hippocampus (Fig. 2A), which is already known as a part of DMN in humans (Greicius et al., 2004), macaques (Mantini et al., 2011; Vincent et al., 2007), and rats (Lu et al., 2012). The ATN (corresponding to Fig. 1B) was strongly functionally connected with caudate and putamen (Fig. 2B). The primary subcortical area connected to the SAN (corresponding to Fig. 1C) was the inferior colliculus (IC) (Fig. 2C). The primary VISs (corresponding to Fig. 1D and 1E) exhibited strong functional connectivities with the lateral geniculate nucleus (LGN), superior colliculus (SC), and ventral lateral (VL), ventral posterior (VP), and pulvinar thalamic nuclei (Fig. 2D). These activations were found in both left and right visual networks. The main subcortical areas in the ORN (corresponding to Fig. 1F) were the ventral striatum, caudate, putamen, and anterior (ANT), laterodorsal (LD), mediodorsal (MD), ventral anterior (VA), and VL thalamic nuclei (Fig. 2E). The main subcortical areas in the higher-order VIS were the SC and LGN for hVIS3 and hVIS4 (Fig. 2F), while there was functional connectivity with the caudate and putamen in these subcortical regions for hVIS1 and hVIS2. In the somatomotor networks, the main subcortical components were the hippocampus and VP thalamic nucleus for the lateral and medial networks (Fig. 2G). In the premotor network, the main subcortical component was VL thalamic nucleus (Fig. 2H). To assign subcortical voxels to networks, the correlation coefficients between the time courses in each cortical network and the time courses in each subcortical voxel were calculated and Fisher’s z-transformed. Then, the network with the highest z-value among all networks was selected and assigned as the main network related to the voxel (Fig. 3).

**Figure 2.**
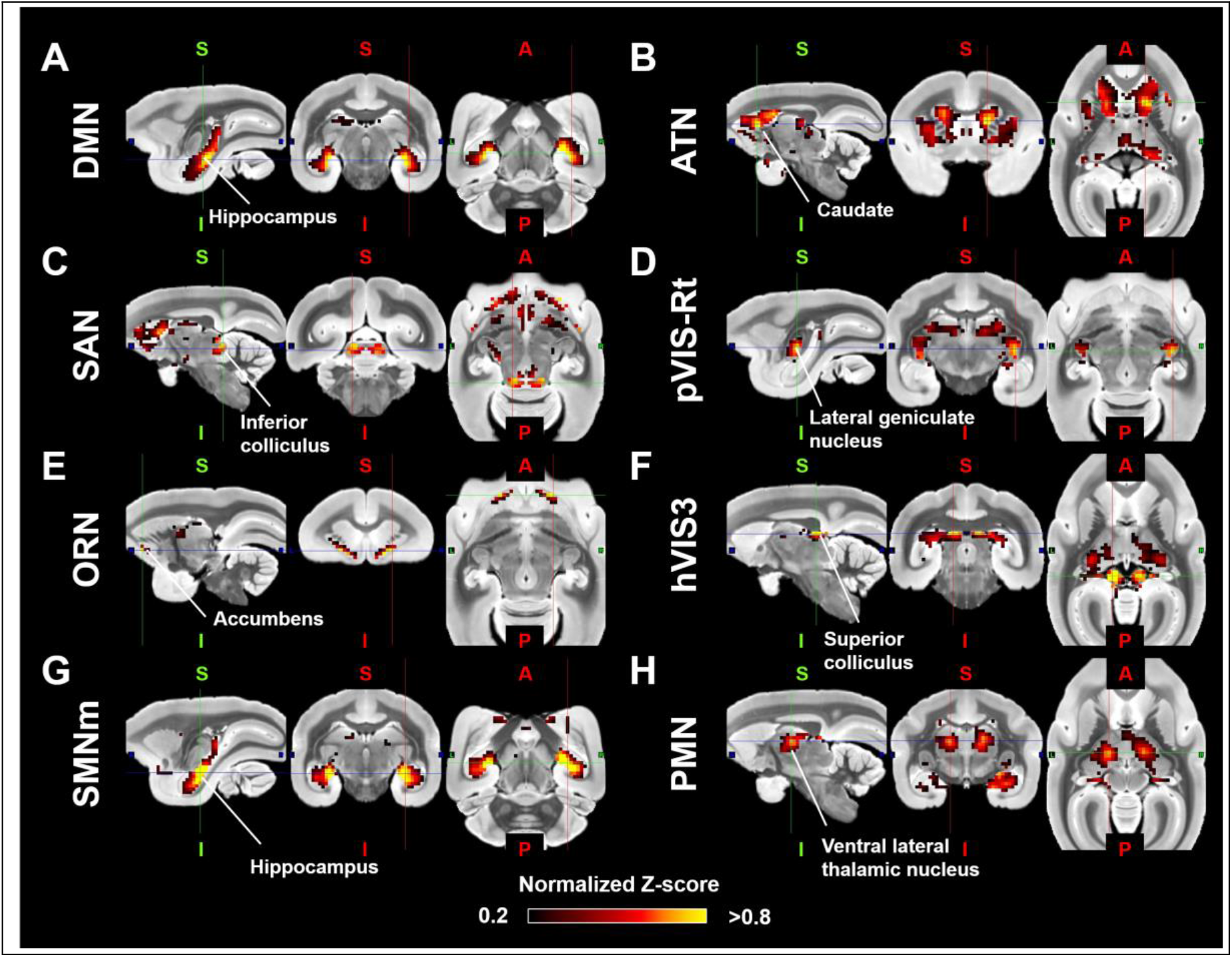
Representative subcortical z-score maps for each resting-state network (RSN). The z-score maps were normalized to be maximum z-value equal to 1, and were shown in sagittal, coronal, and axial slices deemed most representative of the activation patterns. (A) default mode network (DMN: corresponding to Fig. 1A); (B) attention network (ATN: corresponding to Fig. 1B); (C) salience network (SAN: corresponding to Fig. 1C); (D) primary visual network (pVIS: corresponding to Fig. 1E); (E) orbitofrontal network (ORN: corresponding to Fig. 1F); (F) high-order VIS (corresponding to Fig. 1I); (G) somatomotor network (SMN) medial sensory part (corresponding to Fig. 1M); (H) premotor network (PMN) (corresponding to Fig. 1N).

**Figure 3.**
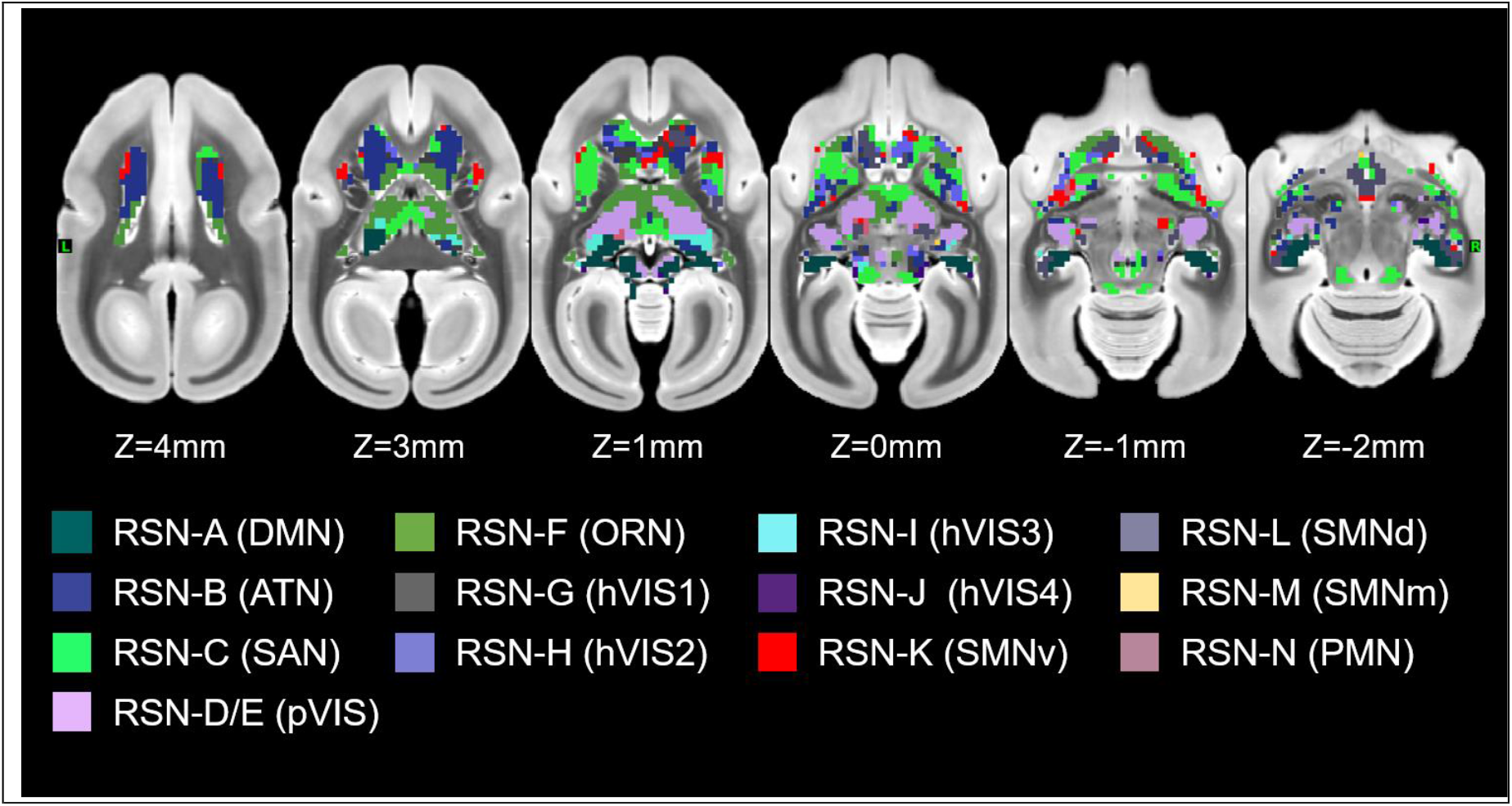
A parcellation of the marmoset subcortical area. The network with the highest z-value among all networks was assigned as the main network related to the voxel. The colors on the surfaces and volumes are corresponding to the name of networks in Figure 1.

### Cortico-subcortical networks in humans

For human RSNs, 10 components were identified as unstructured and/or physiological noise. The remaining 10 components were identified as meaningful functional neural networks. These RSNs were thresholded at z = 3.1 for visual purposes and were named based on the main activation areas with reference to the recent paper, where each cortical partition is assigned to one of the networks (Ji et al., 2019). As such, we identified the DMN (Supplementary Fig. 3A), frontoparietal network (FPN) (Supplementary Fig. 3B), ATN (Supplementary Fig. 3C), two SMNs (ventral: Supplementary Fig. 3D, dorsomedial: 3E), auditory network (AUD) (Supplementary Fig. 3F), two VISs (primary: Supplementary Fig. 3G, high-order: Supplementary Fig. 3H), language network (LAN) (Supplementary Fig. 3I) and cingulo-opercular network (CON) (Supplementary Fig. 3J). Subcortical areas corresponding to each RSN are shown in Supplementary Fig 4. The main subcortical area in the DMN (corresponding to Supplementary Fig. 3A) was the hippocampus (Supplementary Fig. 4A). The FPN (corresponding to Supplementary Fig. 3B) was connected with the caudate and putamen (Supplementary Fig. 4B). The primary subcortical areas connected to the ATN (corresponding to Supplementary Fig. 3C) were the amygdala, SC, and VP and pulvinar thalamic nuclei (Supplementary Fig. 4C). In the SMNs (corresponding to Supplementary Fig. 3D and 3E), the main subcortical components were the hippocampus and VP thalamic nucleus for both ventral and lateral networks (Supplementary Fig. 4D and 4E). The AUD (corresponding to Supplementary Fig. 3F) were functionally connected to all thalamic nuclei (Supplementary Fig. 4F). The primary VIS (corresponding to Supplementary Fig. 3G) exhibited strong functional connectivity with the LGN, SC, and VP and pulvinar thalamic nuclei (Supplementary Fig. 4G), and these activations were also found in the high-order VIS (Supplementary Fig. 3H and 4H). The main subcortical areas in the LAN (corresponding to Supplementary Fig. 3I) were the caudate nucleus and amygdala (Supplementary Fig. 4I). The main subcortical areas in the CON (corresponding to Supplementary Fig. 3J) were the putamen and, ANT, MD, and LD thalamic nuclei (Supplementary Fig. 4J). These subcortical connections in each network were consistent with a previous study (Ji et al., 2019), where they showed caudate, putamen, hippocampus, and amygdala were correlated with FPN, CON, DMN/SMN, and LAN, respectively. The SC and LGN were correlated with primary VIS, which was also consistent with our results. Generally, the subcortical connections except for thalamic nuclei in each human network were similar to the corresponding marmoset networks. For both species, for example, the DMN included hippocampus, and the VIS included LGN and SC. However, thalamic connections in the VIS did not match between marmosets and humans. The VIS in marmosets was strongly connected to the VL thalamic nucleus, while the VIS in humans was mainly connected to the VP and pulvinar thalamic nuclei.

### Comparison of subcortical connectivity profiles

Manhattan distance was used to quantitatively determine how well each subcortical connectivity profile in marmoset RSNs matched the connectivity profile of corresponding human RSNs. Connectivity fingerprints were created for marmosets and humans by determining the mean z-values in seven target regions placed in caudate, putamen, hippocampus, amygdala, SC, IC, and LGN. We did not include the other thalamic VOIs in the fingerprint analysis as these regions are prone to residual global artifacts (Ji et al., 2019). Permutation tests were performed to evaluate statistically significant matches between human and marmoset fingerprints. For each of the ten human RSNs, we tested the hypothesis that the difference between the fingerprints in humans and the target fingerprint in marmosets was smaller than expected by chance. As such, we calculated the Manhattan distance with 10,000 different permutations of the target VOIs in marmosets, following normalization of each fingerprint to a range of 0 (weakest functional connection with any of the target regions) and 1 (strongest functional connection with any of the target regions). A value less than 5 percentile of the histogram of Manhattan distance was considered to be significantly similar fingerprints across species.

The results revealed a number of significant matches between human and marmoset RSNs based on their fingerprints (Fig. 4). The human DMN significantly matched with the marmoset DMN with the strong hippocampus connections (p < 0.05), while the marmoset DMN also exhibited strong FC with the SC (Fig. 5). The FPN in humans had similar subcortical patterns to the ATN, ORN, and VIS1 in marmosets with caudate and putamen connections (p < 0.05; Fig. 6), but the fingerprint of the human ATN did not match with the fingerprint of the marmoset frontoparietal network that we previously labeled ATN in marmosets (Hori et al., 2020b). Instead, in addition to a match with the human FPN, the fingerprint of the marmoset ATN network also matched with the fingerprint of the human LAN (p<0.05; Fig. 7). The primary VIS in humans matched the marmoset primary VIS, high-order VIS3, and VIS4 with strong connections to the SC and LGN (p < 0.05; Fig. 8). The secondary VIS in humans also matched the marmoset high-order VIS (p < 0.05; Fig. 9). The CON in humans matched the marmoset ORN and VIS1 (p < 0.05; Fig. 10).

**Figure 4.**
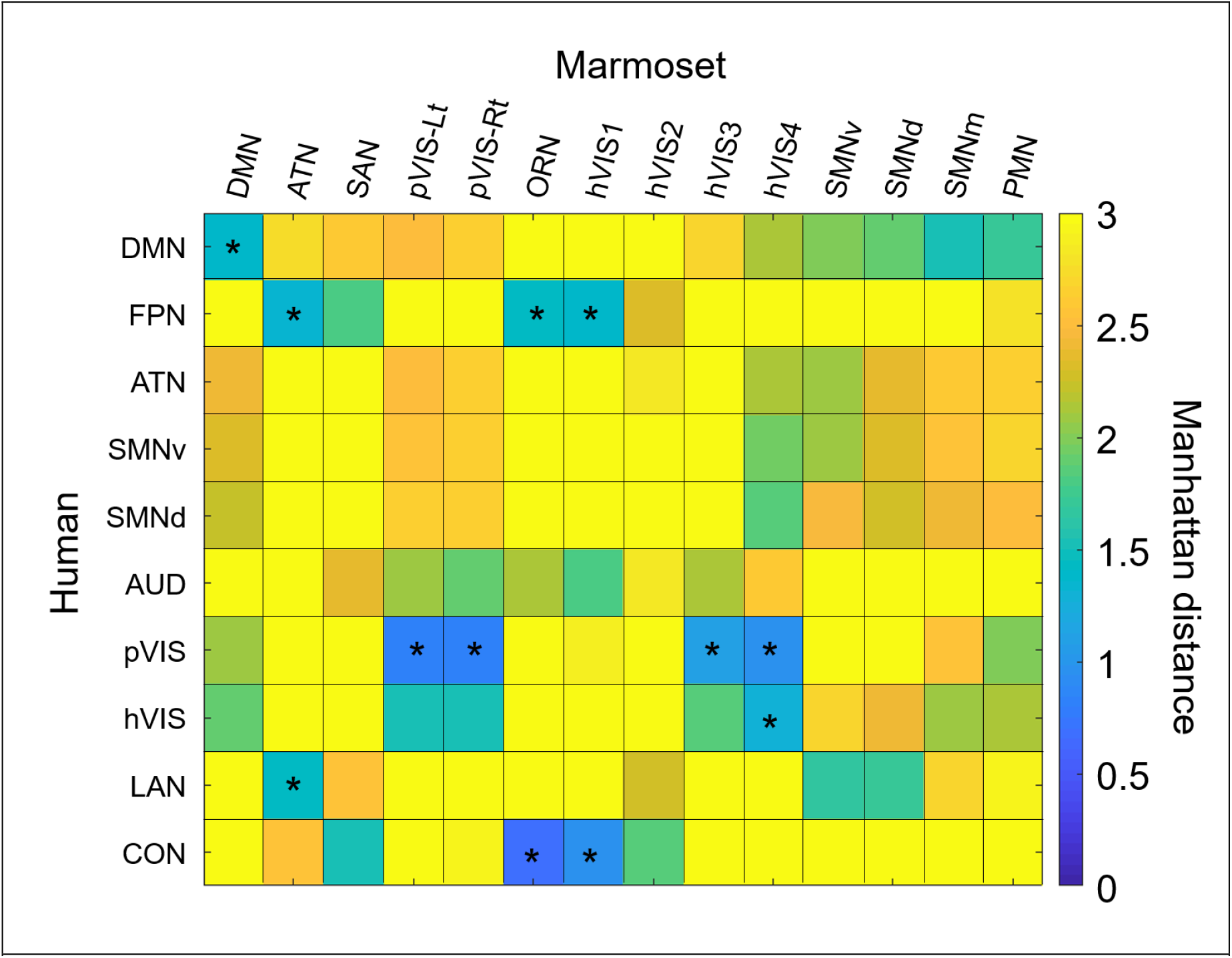
The similarity of subcortical network patterns between marmosets and humans. Manhattan distance between these species were plotted in matrix form. Significant similarities were marked by an asterisk within the matrix.

**Figure 5.**
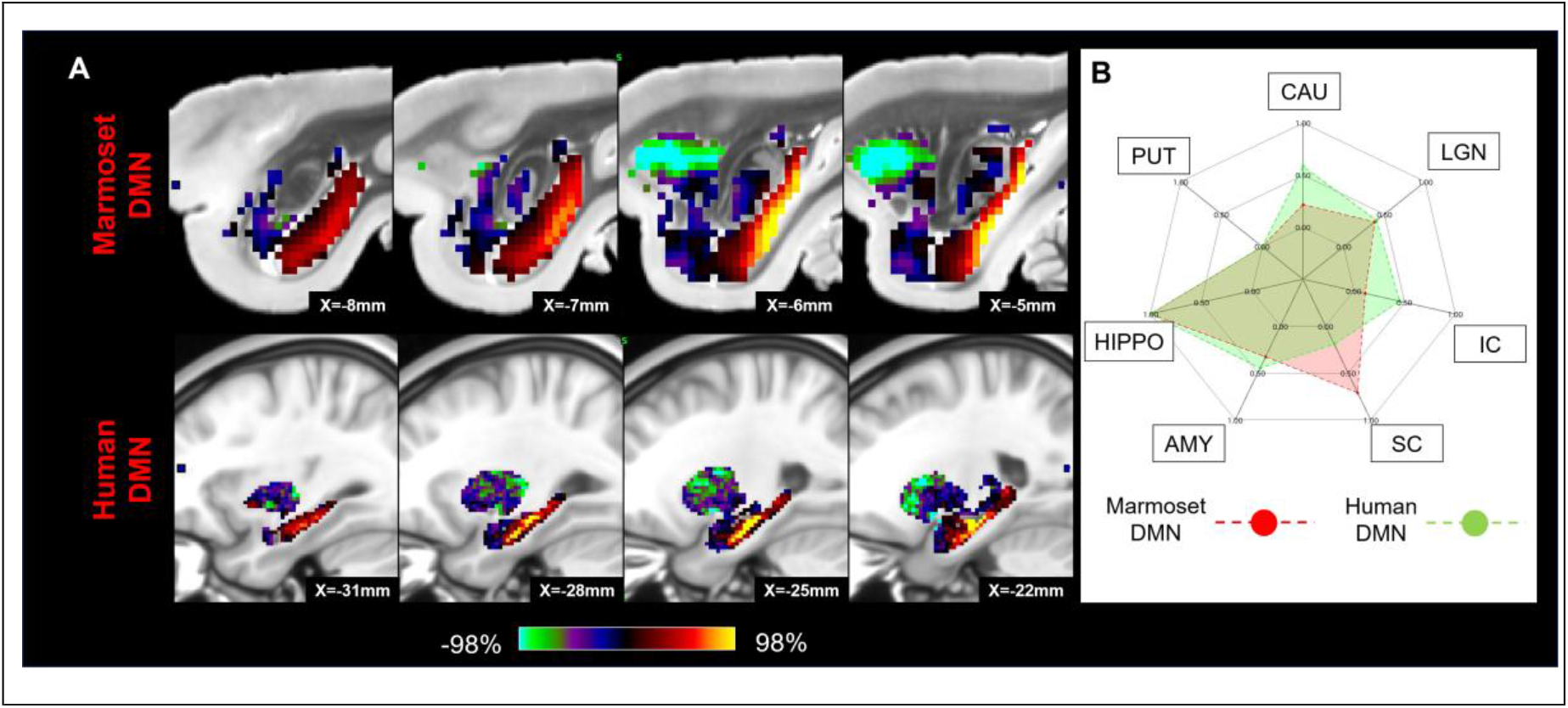
Matching human default mode network (DMN) to marmoset DMN in subcortical areas. (A) Z-score maps were shown in sagittal slices focused on the hippocampus, which has the strongest connections in both species. A single-color palette applies to all two species, but is scaled according to percentile ranges within each species rather than to absolute values. (B) A fingerprint shows the matching connectivity patterns between marmosets and humans. Red and green areas indicate marmoset and human fingerprints, respectively. CAU: caudate; PUT: putamen; HIPPO: hippocampus; AMY: amygdala; SC: superior colliculus; IC: inferior colliculus; LGN: lateral geniculate nucleus.

**Figure 6.**
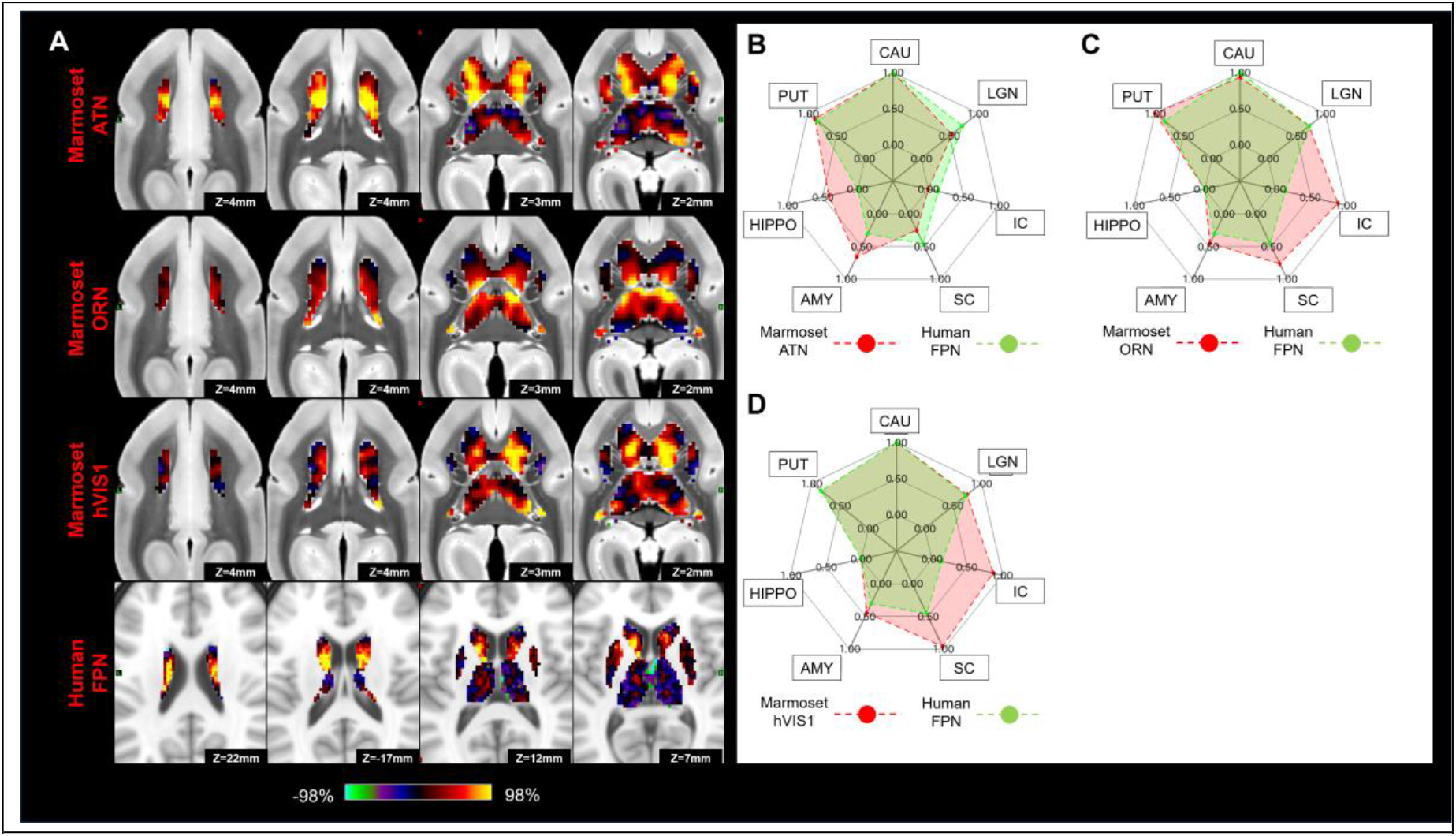
Matching human frontoparietal network (FPN) to marmoset attention (ATN), orbitofrontal (ORN), and high-order visual networks (hVIS1) in subcortical area. (A) Z-score maps were shown in axial slices focused on the caudate, which has the strongest connections in both species. A single-color palette applies to all two species, but is scaled according to percentile ranges within each species rather than to absolute values. (B-D) Fingerprints show the matching connectivity patterns between marmosets and humans. Red and green areas indicate marmoset and human fingerprints, respectively. CAU: caudate; PUT: putamen; HIPPO: hippocampus; AMY: amygdala; SC: superior colliculus; IC: inferior colliculus; LGN: lateral geniculate nucleus.

**Figure 7.**
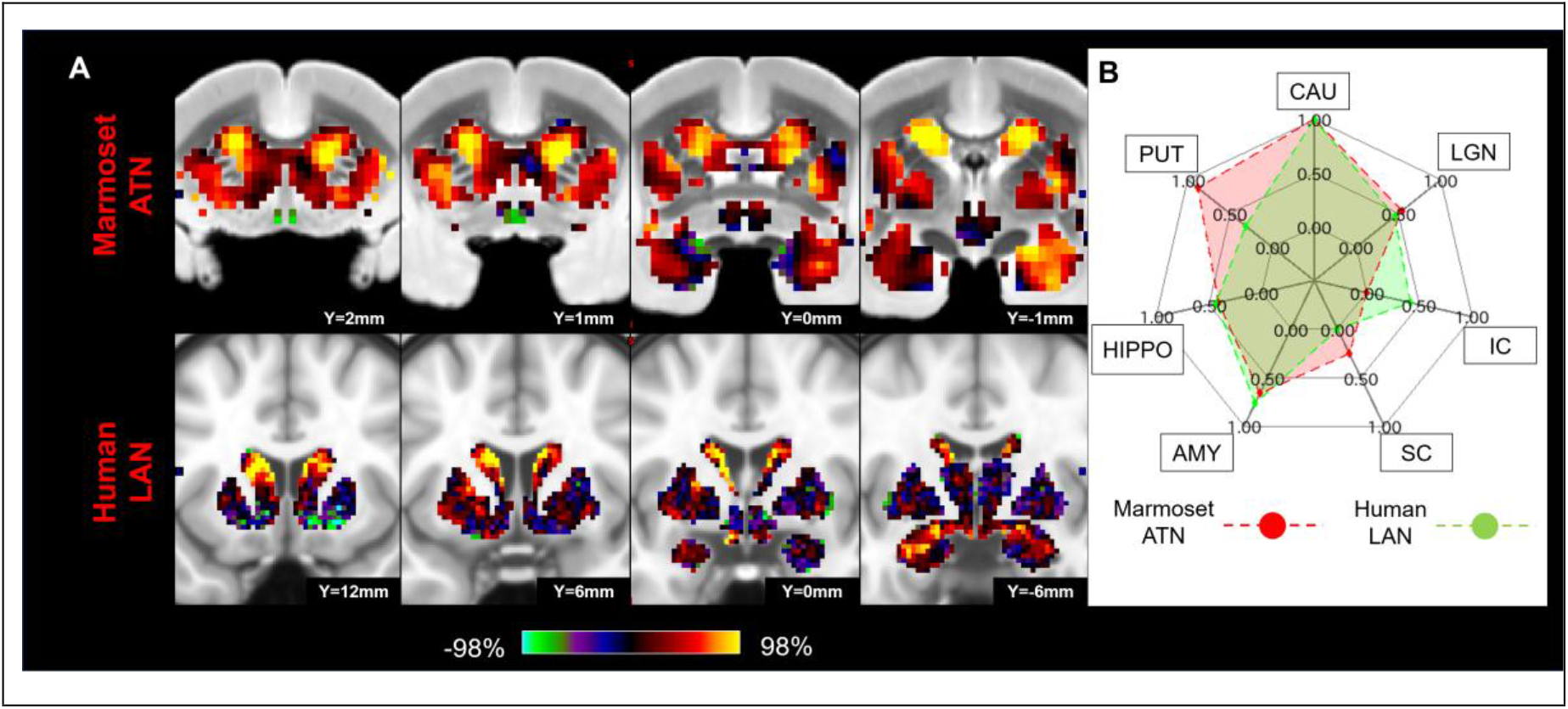
Matching human language network (LAN) to marmoset attention network (ATN) in subcortical area. (A) Z-score maps were shown in coronal slices focused on the caudate and amygdala, which have strong connections in both species. A single-color palette applies to all two species, but is scaled according to percentile ranges within each species rather than to absolute values. (B) Fingerprint shows the matching connectivity pattern between marmosets and humans. Red and green areas indicate marmoset and human fingerprints, respectively. CAU: caudate; PUT: putamen; HIPPO: hippocampus; AMY: amygdala; SC: superior colliculus; IC: inferior colliculus; LGN: lateral geniculate nucleus.

**Figure 8.**
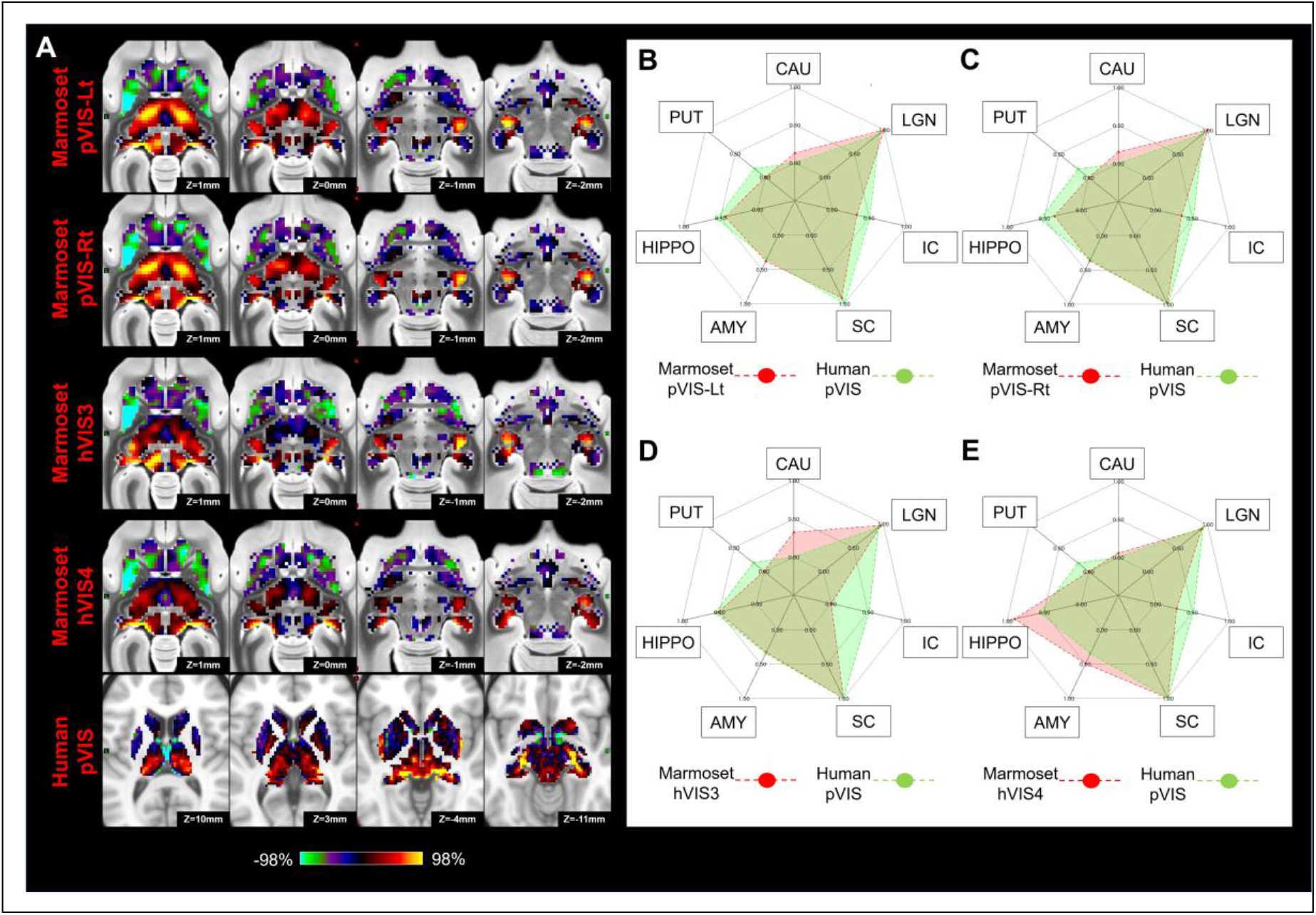
Matching human primary visual network (pVIS) to marmoset VISs (pVIS-Lt, pVIS-Rt, hVIS3, and hVIS4) in subcortical area. (A) Z-score maps for each were shown in axial slices focused on the superior colliculus and lateral geniculate nucleus, which have strong connections in both species. A single-color palette applies to all two species, but is scaled according to percentile ranges within each species rather than to absolute values. (B-E) Fingerprints show the matching connectivity patterns between marmosets and humans. Red and green areas indicate marmoset and human fingerprints, respectively. CAU: caudate; PUT: putamen; HIPPO: hippocampus; AMY: amygdala; SC: superior colliculus; IC: inferior colliculus; LGN: lateral geniculate nucleus.

**Figure 9.**
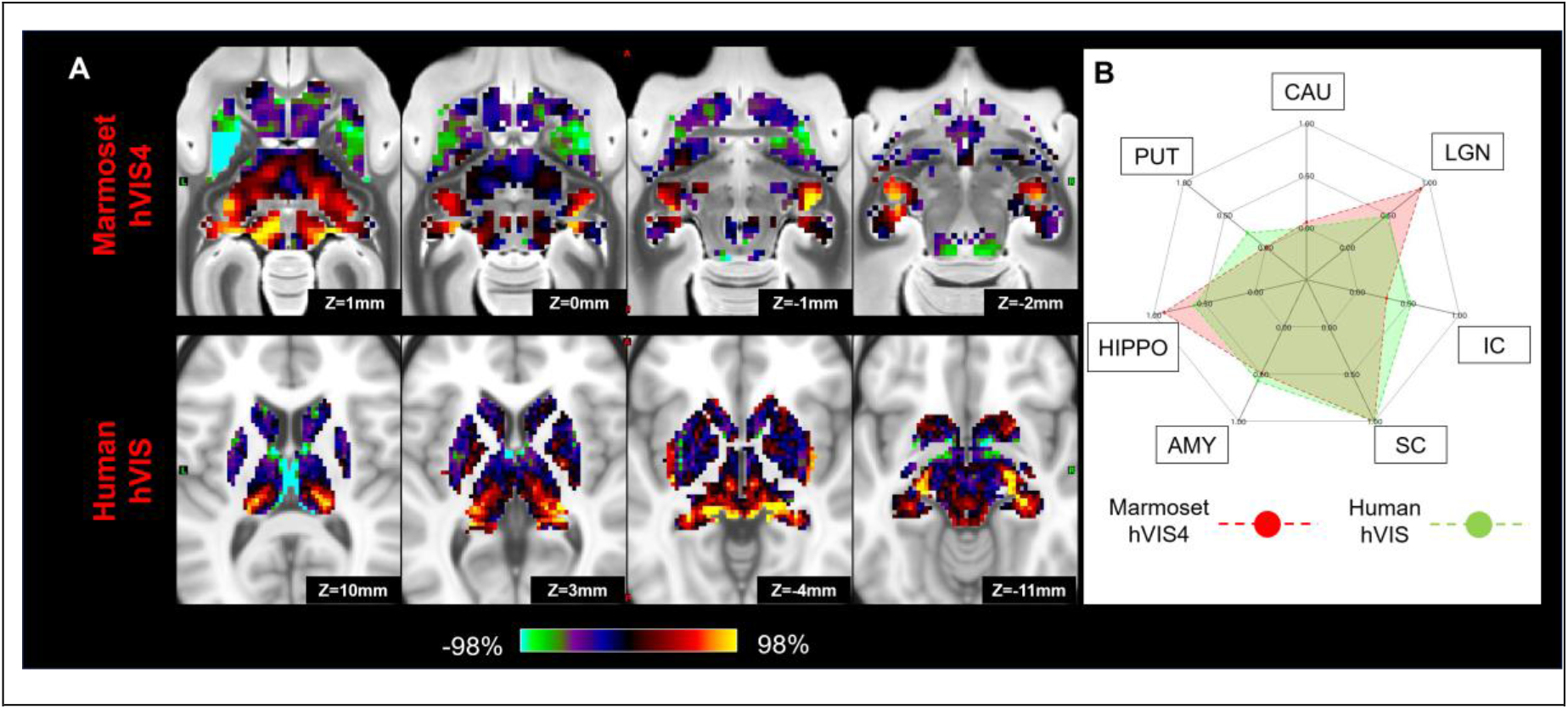
Matching human secondary visual network (hVIS) to marmoset high-order visual network (hVIS4) in subcortical area. (A) Z-score maps were shown in axial slices focused on the superior colliculus and lateral geniculate nucleus, which have strong connections in both species. A single-color palette applies to all two species, but is scaled according to percentile ranges within each species rather than to absolute values. (B) A fingerprint shows the matching connectivity patterns between marmosets and humans. Red and green areas indicate marmoset and human fingerprints, respectively. CAU: caudate; PUT: putamen; HIPPO: hippocampus; AMY: amygdala; SC: superior colliculus; IC: inferior colliculus; LGN: lateral geniculate nucleus.

**Figure 10.**
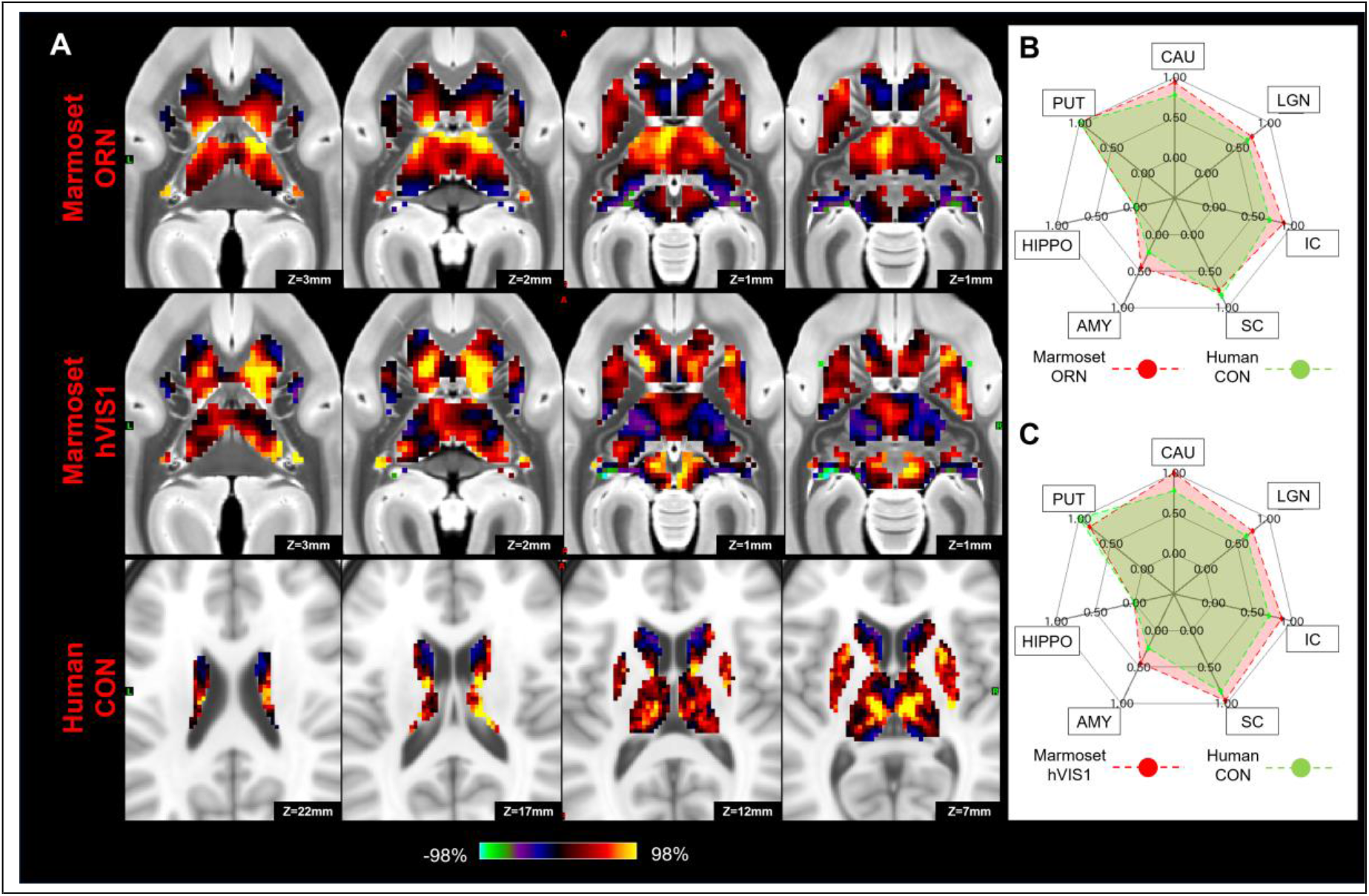
Matching human cingulo-opercular network (CON) to marmoset orbitofrontal (ORN) and high-order visual networks (hVIS1) in subcortical area. (A) Z-score maps were shown in axial slices focused on the caudate and putamen, which have strong connections in both species. A single-color palette applies to all two species, but is scaled according to percentile ranges within each species rather than to absolute values. (B, C) Fingerprints show the matching connectivity patterns between marmosets and humans. Red and green areas indicate marmoset and human fingerprints, respectively. CAU: caudate; PUT: putamen; HIPPO: hippocampus; AMY: amygdala; SC: superior colliculus; IC: inferior colliculus; LGN: lateral geniculate nucleus.

## Discussion

In the present study, we identified the cortico-subcortical functional connections of RSNs in marmosets, then matched these networks with similar human networks based on the fingerprints of their cortico-subcortical functional connectivity profiles. We found that the cortico-subcortical fingerprints of several RSNs matched between marmosets and humans, suggesting a similar functional cortico-subcortical organization of these networks in these two species.

The DMN includes the hippocampus not only in humans (Greicius et al., 2004), but also in macaques (Mantini et al., 2011; Vincent et al., 2007), and rats (Lu et al., 2012). Our results showed that the marmoset DMN includes the hippocampus as well, indicating that it may be a conserved feature across species. Note, however, that the cortical DMN in marmosets was also functionally connected to the SC, which is not the case for the human DMN. In addition to this difference in subcortical connectivity, the marmoset DMN includes a fairly large parietal region that includes areas LIP, VIP, and MIP. Microstimulation in this region around the shallow intraparietal sulcus evokes contralateral saccades (Ghahremani et al., 2019) and single neurons in this region are active for saccadic eye movements (Ma et al., 2020). On the other hand, the parietal region of the human DMN does not include the parietal eye fields defined by multi-modal MRI techniques (Glasser et al., 2016). This discrepancy might produce the difference of the connections with the SC between the species. It also suggests that the marmoset DMN may not have sub serve all of the same functions as the human DMN.

The marmoset ATN consisted of ventral frontal areas (8aV, 45), which are associated with small saccadic eye movements (Selvanayagam et al., 2019). This network was mainly connected to the caudate and putamen, and this subcortical activation pattern was consistent with the human FPN, which includes the FEF and intraparietal areas that are involved in saccade generation (Luna et al., 1998) and attention (Corbetta et al., 1998). Previous human (Raemaekers et al., 2006; Raemaekers et al., 2002) and macaque studies (Hikosaka and Wurtz, 1989; Phillips and Everling, 2012) have shown activations in the striatum (both caudate and putamen) during saccade tasks. The parietal component of the marmoset ATN, however, lies anterior to area LIP which is activated by saccadic eye movements (Schaeffer et al., 2019d) and where saccades can be evoked by electrical microstimulation in marmosets (Ghahremani et al., 2019), arguing perhaps against a pure role of this network in saccadic eye movements. In fact, we found that the human language network (LAN) also matched the marmoset ATN in terms of its cortico-subcortical connectivity fingerprint. In both species, the networks showed strong functional connectivity with the caudate. In addition, there are also clear similarities in the cortical regions between the marmoset ATN and the human LAN. Broca’s area (area 44, 45) is a prominent part of the human LAN and the marmoset ATN network also includes area 45. Single neurons in marmoset area 45 and 8aV respond to marmoset vocalizations and many are active for vocalizations (Miller et al., 2015), supporting a role of this area in vocalization. The finding that the cortico-subcortical fingerprint of the ATN matched with both the human FPN and the LAN may suggest that this core frontoparietal network is the evolutionary precursor to these networks. Although we have labeled this network as ATN here to be consistent with a previous paper (Hori et al., 2020b), a better label would probably be just FPN for this network, consistent with an older report from our lab that used ICA to identify RSNs in anesthetized marmosets (Ghahremani et al., 2016).

The subcortical pattern in the human FPN also showed a match to the marmoset hVIS1. The main cortical activations in hVIS1 were along with the ventral visual stream including TE3, V4T, and FST (Hung et al., 2015a; Schaeffer et al., 2019d). This is consistent with human and macaque studies showing that the FPN includes a part of ventral visual stream (Hutchison et al., 2012; Ji et al., 2019; Thomas Yeo et al., 2011). In addition, both human and macaque FEF are functionally connected to these regions (Hutchison et al., 2012). As such, the hVIS1 in marmosets seems to correspond to the temporal regions in the human FPN. Interestingly, the subcortical pattern in the hVIS1 also showed a match to the human CON as well as FPN. These two functional networks display increased activity during the performance of complex cognitive tasks (Dosenbach et al., 2006; Sheffield et al., 2015; Wallis et al., 2015), and both are associated with top-down control associated with executive functioning (Dosenbach et al., 2007). Taken together with our findings, the marmoset hVIS1 might be related to both FPN and CON through the putamen and caudate and play an important role in top-down cognitive processing.

The cortical visual networks in marmosets were strongly connected to the SC, LGN, VP, and PUL. These regions are known to be associated with the visual system (Hung et al., 2015a; Hung et al., 2015b), and are structurally connected to visual-related cortices (Kaas and Lyon, 2007; Solomon and Rosa, 2014; Zeater et al., 2019). We found that these subcortical activation patterns in marmoset corresponded well to those in humans, suggesting that the visual systems have a similar cortico-subcortical organization in both species. Previous anatomical studies and electrophysiological recordings in marmosets have also shown that this species’ cortical visual hierarchy closely resembles that of other primates, including humans (McDonald et al., 2014; Mitchell and Leopold, 2015; Yu and Rosa, 2010).

## Conclusion

We have shown here that many of the marmoset RNSs can be matched to human RSNs based on their cortico-subcortical fingerprint. While this suggests a similar cortico-subcortical network organization in marmosets and humans, our results also show that there are differences in the connectivity profiles that likely have consequences on the actual functions of these RSNs. Electrophysiological and task-based fMRI studies in marmosets will be necessary to further investigate functional similarities and differences in RSN organization between the two species.

## Methods

### Animal preparation

All surgical and experimental procedures were in accordance with the Canadian Council of Animal Care policy and a protocol approved by the Animal Care Committee of the University of Western Ontario Council on Animal Care. All animal experiments complied with the Animal Research: Reporting *In Vivo* Experiments (ARRIVE) guidelines. Four male common marmosets weighting 390 g (3 years old), 245 g (1.5 years old), 330g (1.5 years old), and 360g (1.5 years old), and one female marmoset weighting 306 g (1.3 years old) were used in *in-vivo* and *ex-vivo* study, respectively.

Four marmosets for *in-vivo* experiments underwent surgery to implant a head chamber to fix the head during MRI acquisition as described in previous reports (Johnston et al., 2018; Schaeffer et al., 2019c). Briefly, the marmoset was placed in a stereotactic frame (Narishige Model SR-6C-HT), and several coats of adhesive resin (All-bond Universal Bisco, Schaumburg, Illinois, USA) were applied using a microbrush, air dried, and cured with an ultraviolet dental curing light. Then, a dental cement (C & B Cement, Bisco, Schaumburg, Illinois, USA) was applied to the skull and to the bottom of the chamber, which was then lowered onto the skull via a stereotactic manipulator to ensure correct location and orientation. The chamber was 3D printed at 0.25 mm resolution using stereolithography and a clear photopolymer resin (Clear-Resin V4; Form 2, Formlabs, Somerville, Massachusetts, USA). The marmosets were first acclimatized to the animal holder, head fixation system, and a mock MRI environment prior to the first imaging session (Silva et al., 2011). Each marmoset was trained over the course of three weeks. During the first week, marmosets entered the tube and were constrained using only the neck and tail plates for increasingly long periods of time (up to 30 minutes). During the second week, the restraint tube was inserted into a mock MRI tube (a 12 cm inner diameter tube) to simulate the scanner environment; MRI sounds were played at increasingly loud volumes (up to 80 dB) for increasingly long durations, up to 60 minutes sessions. In week 3, marmosets were head-fixed via the fixation pins, inserted into the mock MRI tube and exposed to the MRI sounds. Within each session, the animals are presented with reward items (pudding or marshmallow fluff) for remaining still (calmly facing forward, with minimal movement of limbs). Throughout the training sessions, the behavioral rating scale described by Silva et al. (2011) was used to assess the animals’ tolerance to the acclimatization procedure by the end of week 3, all three marmosets scored 1 or 2 on this assessment scale (Silva et al., 2011), showing calm and quiet behavior, with little signs of agitation.

To create volumes of interests (VOIs) in subcortical area, we acquired *ex-vivo* MRI as it allowed for longer scanning time at a much higher resolution (0.1 mm isotropic). To prepare for *ex-vivo* MRI, one marmoset was euthanized through transcardial perfusion and its brain was extracted at the end of the procedure. Anesthesia was initially induced with 30 mg/kg of ketamine and maintained with 4% isoflurane in 1.5-2% oxygen. The animal was then transcardially perfused with 0.9% sodium chloride solution, followed by 10% formaldehyde buffered solution (formalin). The brain was then extracted and stored in 10% buffered formalin for over a week.

### Image acquisition

For the *in-vivo* experiment, each animal was fixed to the animal holder using a neck plate and a tail plate. The animal was then head-fixed using fixation pins in the MRI room to minimize the time in which the awake animal was head fixed (Schaeffer et al., 2019c). Once fixed, a lubricating gel (MUKO SM321N, Canadian Custom Packaging Company, Toronto, Ontario, Canada) was squeezed into the chamber and applied to the brow ridge to reduce magnetic susceptibility.

Data were acquired using a 9.4 T 31 cm horizontal bore magnet (Varian/Agilent, Yarnton, UK) and Bruker BioSpec Avance III HD console with the software package Paravision-6 (Bruker BioSpin Corp, Billerica, MA), a custom-built high-performance 15-cm-diameter gradient coil with 400-mT/m maximum gradient strength (Handler et al., 2020), and the 5-channel receive coil (Schaeffer et al., 2019c). Radiofrequency transmission was accomplished with a quadrature birdcage coil (12-cm inner diameter) built in-house. All imaging was performed at the Centre for Functional and Metabolic Mapping at the University of Western Ontario.

Functional images were acquired with 6-22 functional runs (at 400 or 600 volumes each) for each animal in the awake condition, using gradient-echo based single-shot echo-planar imaging sequence with the following parameters: TR = 1500 ms, TE = 15 ms, flip angle = 40°, field of view (FOV) = 64 × 64 mm, matrix size 128 × 128, voxel size 0.5 mm isotropic, slices = 42, bandwidth = 500 kHz, generalized autocalibrating parallel acquisition (GRAPPA) acceleration factor (anterior-posterior) = 2. Total scan time for all functional imaging was ~14h. A T2-wighted image (T2w) was also acquired for each animal using rapid imaging with refocused echoes (RARE) sequences with the following parameters: TR = 5500 ms, TE = 53 ms, FOV = 51.2 × 51.2 mm, matrix size = 384 × 384, voxel size = 0.133 × 0.133 × 0.5 mm, slice 42, bandwidth = 50 kHz, GRAPPA acceleration factor (anterior-posterior) = 2.

For *ex-vivo* imaging, a formalin-fixed marmoset brain was submerged in lubricant (Christo-lube; Lubrication technology Inc., Franklin Furnace, OH) to avoid magnetic susceptibility-related distortion artifacts, and three-dimensional multi-echo spin-echo images were acquired as following parameters: TR = 200 ms, TE = 3. 5, 8.5, 13.5, 18.5, 23.5 ms, FOV = 33 × 28.8 × 36 mm, matrix size = 330 × 288 × 360, voxel size = 0.1 mm isotropic resolution, average=4. The average image across different TE images was calculated to increase the signal-to-noise ratio (SNR) and it was used to create the subcortical volume-of-interests (VOIs).

### Marmoset image preprocessing

Data was preprocessed using FSL software (Smith et al., 2004). Raw MRI images were first converted to Neuro Informatics Technology Initiative (NIfTI) format (Li et al., 2016) and reoriented from sphinx position. Brain masks for *in-vivo* images were created using FSL tools and the National Institutes of Health (NIH) T2w brain template (Liu et al., 2018) For each animal, the brain-skull boundary was first roughly identified from individual T2w using the brain extraction tool (BET) with the following options; radius of 25-40 and fractional intensity threshold of 0.3 (Smith, 2002). Then, the NIH T2w brain template was linearly and non-linearly registered to the individual brain image using FMRIB’s linear registration tool (FLIRT) and FMRIB’s nonlinear registration tool (FNIRT) to more accurately create the brain mask. After that, the brain was extracted using the brain mask. RS-fMRI images were corrected for motion using FLIRT. Principal component analysis (PCA) was applied to remove the unstructured noise from the RS-MRI time course, followed by independent component analysis (ICA) with the decomposition number of 200 using Multivariate Exploratory Linear Optimized Decomposition into the Independent Components (MELODIC) module of the FSL software package. Obtained components were classified as signal or noise (such as eye movement, CSF pulsation, heart rate, and respiratory artifacts) based on the criteria as shown a previous report (Griffanti et al., 2017), and noise components were regressed out from the rfMRI time course using FSL tool (fsl_regfilt). All rfMRI images were finally normalized to the NIH template using rfMRI-to-T2w and T2w-to-template transformation matrices obtained by FLIRT and FNIRT, followed by spatial smoothing by Gaussian kernel with the full width of half maximum value of 1.0 mm. The *ex-vivo* structure image was also normalized to the NIH template using FLIRT and FNIRT.

### Cortico-subcortical functional networks in marmosets

The group ICA analysis was first implemented for only cortical area 10 times with different dimension numbers (from 16 to 25) to identify optimal dimensionality using MELODIC module of the FSL software package – the 20 components solution was selected to be an appropriate representative of meaningful components with reference to previous reports of marmoset functional networks (Belcher et al., 2013; Ghahremani et al., 2016; Hori et al., 2019b). Second, a spatial regression approach was used to obtain the temporal dynamics for each cortical component within each scan’s fMRI data sets (Filippini et al., 2009). In this process, the full set of group-ICA spatial templates were used in a linear model fit against the separate fMRI data sets. Finally, we calculated correlation coefficients between the time courses in each cortical network and the time courses in each subcortical voxel using FSL’s FEAT. The functional connectivity maps (z-score maps) in the subcortical areas were then averaged across scans. To assign each subcortical voxel to one of the networks, one network having the highest z-value was assigned in each voxel.

### Cortico-subcortical functional networks in humans

Human connectome project (HCP) datasets were used for human analysis (Van Essen et al., 2013). RS-fMRI data for 100 subjects (4 scans for each subject, namely total 400 scans) preprocessed with the HCP functional pipeline, including motion correction, distortion correction, normalization to Montreal Neurological Institute (MNI) template space, and FMRIB’s ICA-based X-noiseifier (FIX) denoising (Salimi-Khorshidi et al., 2014) were downloaded from the HCP website (https://www.humanconnectome.org/). Group ICA analysis was performed for only cortical areas with 20 dimensions. After that, temporal dynamics for each cortical component within each scan’s data were obtained by a spatial regression approach using group ICA templates, and correlation coefficients between the time courses in each cortical network and the time courses in each subcortical voxel were calculated in the same way as in the marmoset analysis. Finally, the functional connectivity maps (z-score maps) in the subcortical areas were then averaged across scans.

### Subcortical volume of interest

To identify the subcortical areas associated with each marmoset network, we applied the subcortical atlas supplied by the NIH Marmoset Brain template (Liu et al., 2018), where the thalamus is not parcellated into subthalamic nuclei. Based on the ex-vivo image normalized to the NIH template, we created the subthalamic VOIs (anterior (AN), laterodorsal (LD), mediodorsal (MD), ventral anterior (VA), ventral lateral (VL), ventral posterior (VP) and pulvinar) with reference to the Paxinos atlas (Paxinos et al., 2012). For human subcortical VOIs, standard mesh atlas for subcortical area supplied by HCP pipeline was used (Atlas_ROIs_2.nii.gz), which does not have VOIs of subthalamic nuclei, lateral geniculate nucleus (LGN), superior colliculus (SC) and inferior colliculus (IC). For thalamic nuclei, the histological-based atlas supplied by NeuroImaging and Surgical Technologies Lab was used (Xiao et al., 2015; Xiao et al., 2012). Also, a radiologist made the VOIs for LGN, SC and IC based on the MNI T1w template.

### Comparison of subcortical connectivity profiles

To determine whether the subcortical connectivity profiles are either similar or dissimilar from each other, we used Manhattan distance among each connectivity fingerprint (Mars et al., 2018; Mars et al., 2016). Connectivity fingerprints were created for marmosets and humans by determining the mean z-values in seven target regions placed in the caudate, putamen, hippocampus, amygdala, SC, IC, and LGN. We normalized the fingerprint to a range between 0 (weakest connection with any of the target areas) and 1 (strongest connection with any of the target areas) to compare a pattern of connections with target areas, rather than absolute strength. Permutation testing was used to test the significance of the match between each of the marmoset and human subcortical network by calculating 10,000 different permutations of the fingerprint target networks in marmosets. p < 0.05 is considered as significantly smaller Manhattan distance than expected chance. This analysis was performed using custom tools written in Matlab (the Mathworks, Natick, MA, USA).

## Acknowledgements

This work was supported by the Canadian Institutes of Health Research (FRN 148365, FRN 353372) and the Canada First Research Excellence Fund to BrainsCAN. Human data were provided by the Washington University-University of Minnesota Consortium of the Human Connectome Project (WU-Minn HCP; Principal Investigators: David Van Essen and Kamil Ugurbil; 1U54MH091657) funded by the 16 NIH Institutes and Centers that support the NIH Blueprint for Neuroscience Research; and by the McDonnell Center for Systems Neuroscience at Washington University. We also thank Miranda Bellyou for animal preparation and care and Dr. Alex Li for scanning assistance.

## Supplementary Figure

**Supplementary Figure 1.**
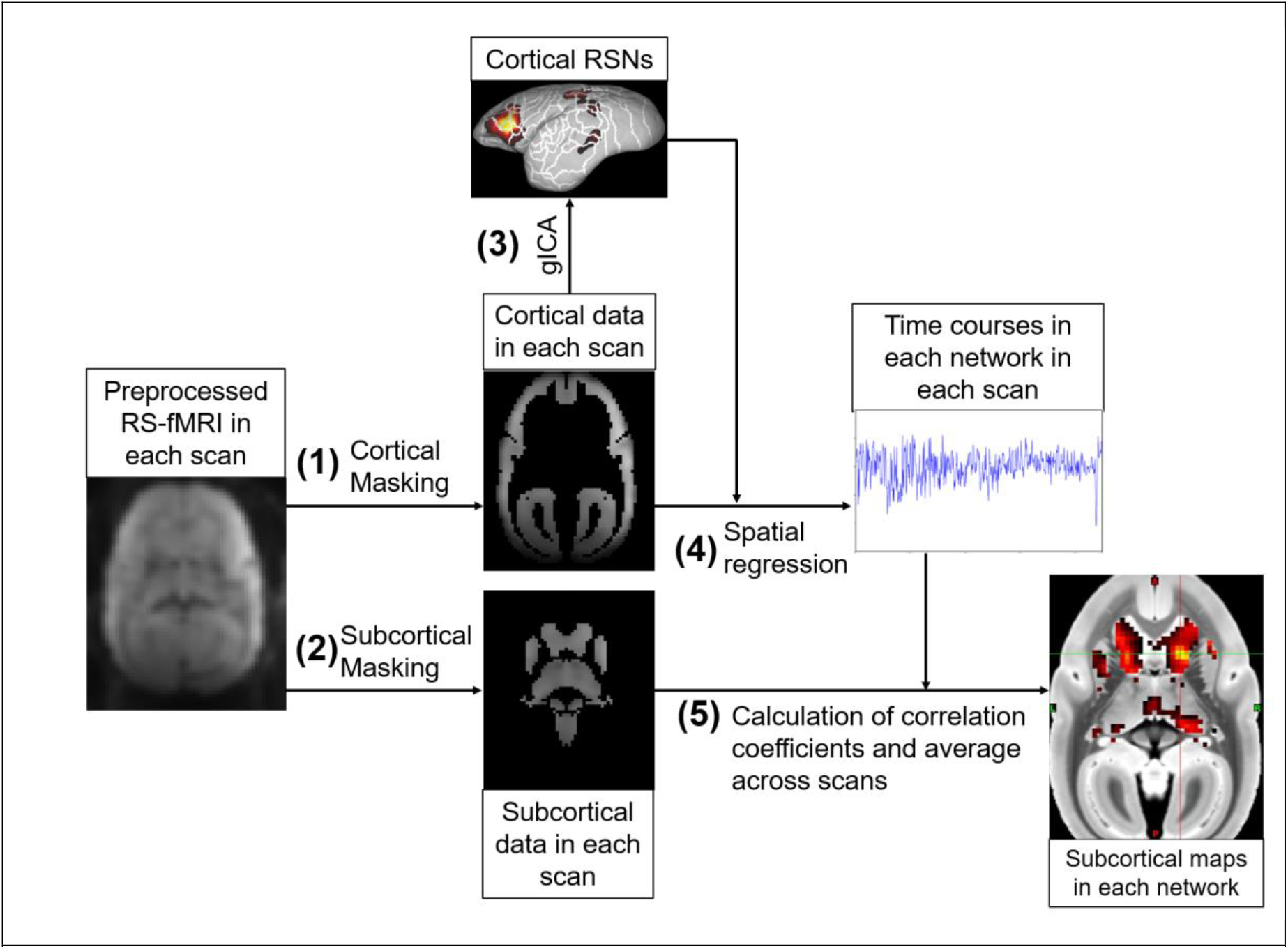
Flow chart of analysis to calculate the subcortical connectivity maps. Each RS-fMRI scan was preprocessed, and (1) cortical and (2) subcortical regions were extracted using masks. (3) Using all cortical RS-fMRI datasets, group ICA (gICA) was performed so that 14 and 10 cortical resting-state networks (RSNs) were identified for marmosets and humans, respectively. (4) The time courses of each network in each scan were calculated using spatial regression technique and obtained cortical RSNs, (5) then correlation coefficients between the time courses in each cortical network and the time courses in each subcortical voxel were calculated. Obtained correlation coefficient maps in each network were averaged across scans.

**Supplementary Figure2.**
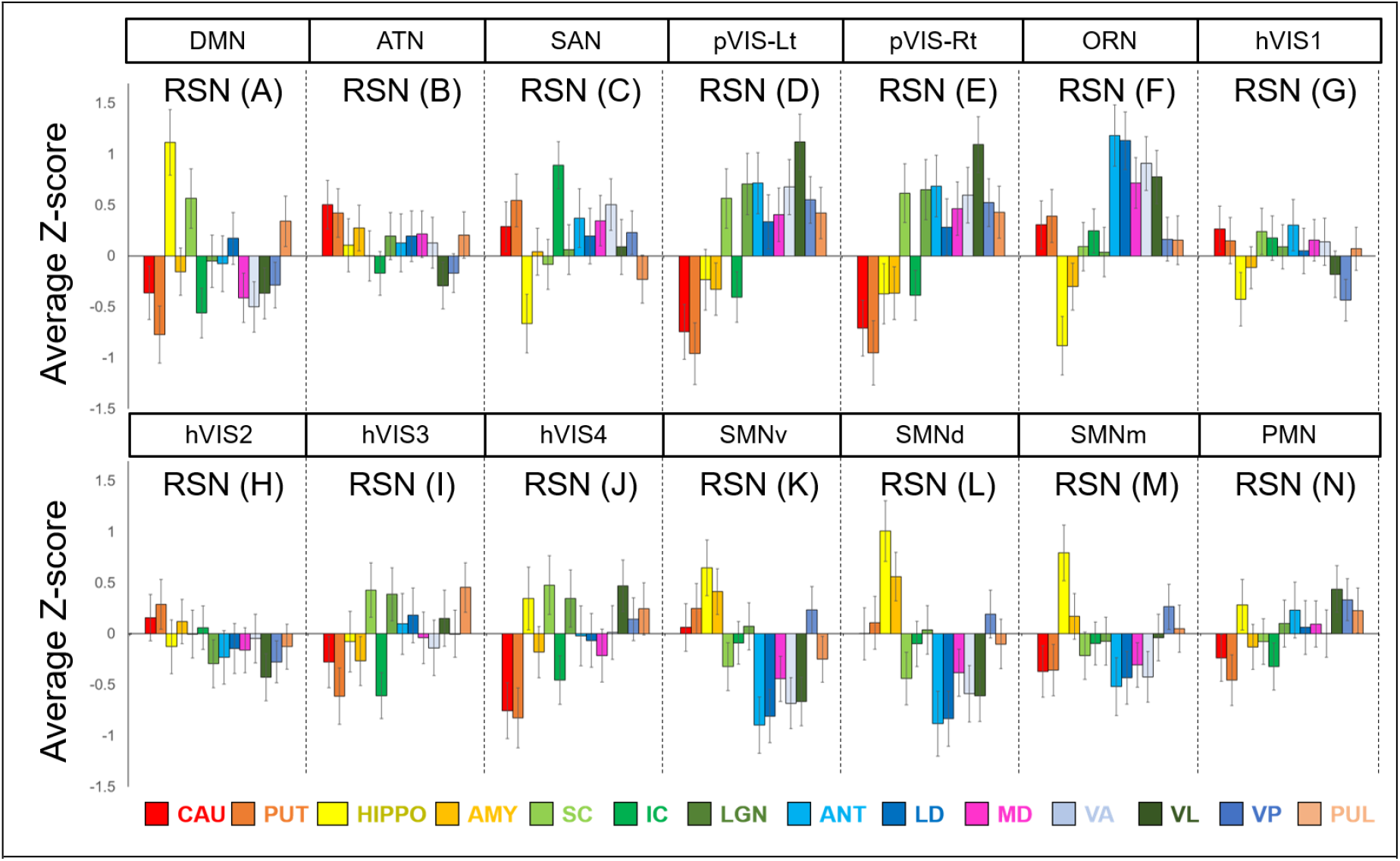
Mean z-score values in each subcortical area in each marmoset resting-state network (RSN). The error bars represent the standard error of the mean. The RSNs described here are corresponding to those in Fig. 1. CAU: caudate; PUT: putamen; HIPPO: hippocampus; AMY: amygdala; SC: superior colliculus; IC: inferior colliculus; LGN: lateral geniculate nucleus; ANT: anterior part of thalamic nucleus; LD: laterodorsal thalamic nucleus; MD: mediodorsal thalamic nucleus; VA: ventral anterior thalamic nucleus; VL: ventral lateral thalamic nucleus; VP: ventral posterior thalamic nucleus; PUL: pulvinar nucleus.

**Supplementary Figure 3.**
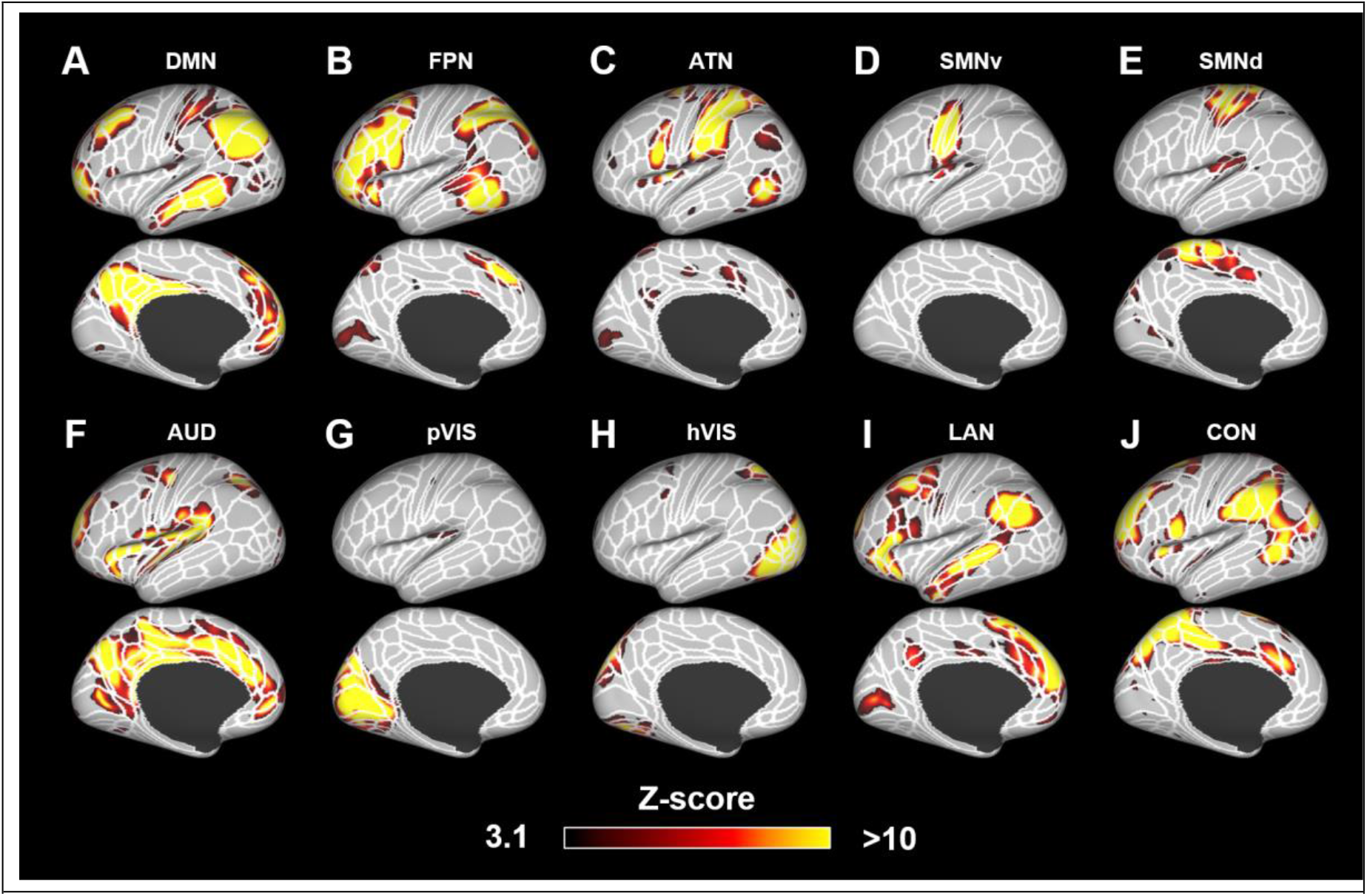
Ten components identified as resting-state networks in humans. These networks were labeled based on a previous study (Ji et. al., 2019) as follows: (A) default mode network (DMN); (B) frontoparietal network (FPN); (C) attention network (ATN); (D) somatomotor network ventral part (SMN1); (E) SMN dorsomedial part (SMN2); (F) auditory network (AUD); (G) primary visual network (pVIS); (H) high-order VIS (hVIS); (I) language network (LAN); (J) cingulo-opercular network (CON). Color bar represents the z-score of these correlation patterns thresholding at 3.1. White lines show the parcellation borders created based on the multimodal magnetic resonance images from Human Connectome Project (Glasser et al., 2016).

**Supplementary Figure 4.**
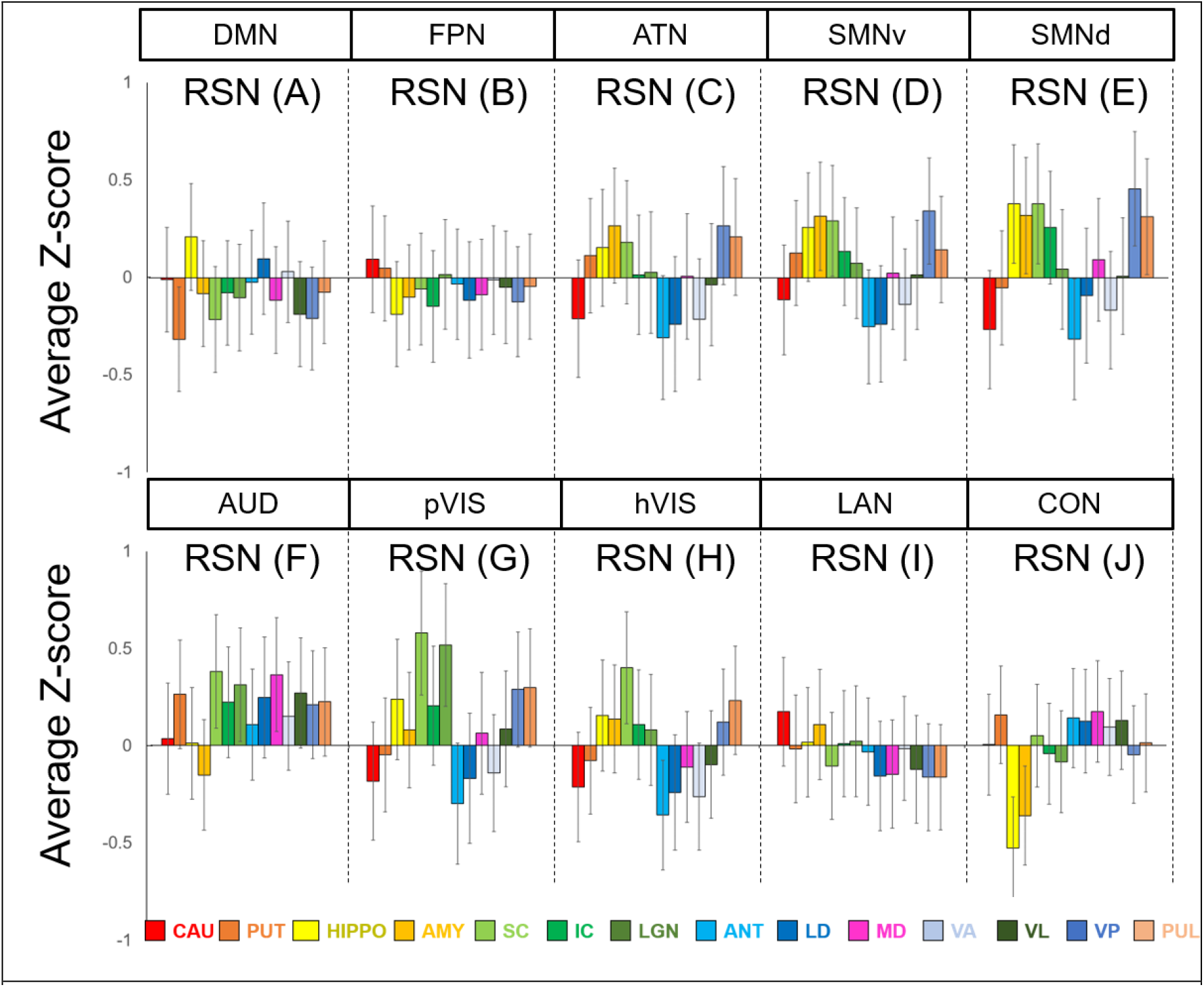
Mean z-score values in each subcortical area in each human resting-state network (RSN). The error bars represent the standard error of the mean. The RSNs described here are corresponding to those in Supplementary fig. 2. CAU: caudate; PUT: putamen; HIPPO: hippocampus; AMY: amygdala; SC: superior colliculus; IC: inferior colliculus; LGN: lateral geniculate nucleus; ANT: anterior part of thalamic nucleus; LD: laterodorsal thalamic nucleus; MD: mediodorsal thalamic nucleus; VA: ventral anterior thalamic nucleus; VL: ventral lateral thalamic nucleus; VP: ventral posterior thalamic nucleus; PUL: pulvinar nucleus.

